# Shifts in neural tuning systematically alter sensorimotor learning ability

**DOI:** 10.1101/2024.07.29.605715

**Authors:** Takuji Hayashi, Ken Takiyama, Maurice A. Smith, Daichi Nozaki

## Abstract

Sensorimotor learning can change the tuning of neurons in motor-related brain areas and rotate their preferred directions (PDs). These PD rotations are commonly interpreted as reflecting motor command changes; however, cortical neurons that display PD rotations also contribute to sensorimotor learning. Sensorimotor learning should, therefore, alter not only motor commands but also the tuning of neurons responsible for this learning, and thus impact subsequent learning ability. Here, we investigate this possibility with computational modeling and by directly measuring adaptive responses during sensorimotor learning in humans. Modeling shows that the PD rotations induced by sensorimotor learning, predict specific anisotropic changes in PD distributions that in turn predict a specific spatial pattern of changes in learning ability. Remarkably, experiments in humans then reveal large, systematic changes in learning ability in a spatial pattern that precisely reflects these model-predicted changes. We find that this pattern defies conventional wisdom and implements Newton’s method, a learning rule where the step size is inversely proportional rather than proportional to the learning gradient’s amplitude, limiting overshooting in the adaptive response. Our findings indicate that PD rotation provides a mechanism whereby the motor system can simultaneously learn how to move and learn how to learn.

## INTRODUCTION

Neurons in motor-related brain areas typically exhibit activity modulated by several properties of reaching movements, perhaps the strongest of which is the direction of movement (Georgopoulos et al., 1986, 1982; Kalaska et al., 1989). This tuning function, characterized by the preferred direction (PD) where a neuron’s activity is maximal, provides invaluable information not only for understanding how movement is represented in the brain (Georgopoulos et al., 1986, 1982; Kalaska et al., 1989) but also for designing controllers for brain-computer interfaces (Hochberg et al., 2012, 2006; Taylor et al., 2002). Critically, a number of studies have demonstrated plasticity in neural PDs, as motor learning arising from both kinematic (Alexander and Crutcher, 1990a, 1990b; Shen and Alexander, 1997) and dynamic (Gandolfo et al., 2000; Li et al., 2001; Perich and Miller, 2017; Rokni et al., 2007) perturbations can result in systematic rotations of these PDs.

PD rotation is thought to reflect changes in the motor commands resulting from motor learning (Hirashima and Nozaki, 2012; Rokni et al., 2007; Takiyama and Okada, 2012). However, it is also widely believed that neurons in motor cortical areas that display PD rotation, can serve as neuronal learning units (or motor learning primitives) (Donchin et al., 2003; Herzfeld et al., 2014; Hwang et al., 2003; Rokni et al., 2007; Sing et al., 2009; Thoroughman and Shadmehr, 2000; Wainscott et al., 2005; Yokoi et al., 2011). If this were the case, then the changes in neural tuning characterized by PD rotation would also alter the process of learning. We, therefore, hypothesize that PD rotation does not merely occur as a consequence of motor learning, but also determines subsequent motor learning ability. A network model based on this framework incorporates neural units with plastic output weights that are recruited according to tuning functions that change with PD rotation. Previous models have been based on units with plastic output weights, but fixed tuning functions unaltered by learning (Donchin et al., 2003; Herzfeld et al., 2014; Hwang et al., 2003; Sing et al., 2009; Thoroughman and Shadmehr, 2000; Wainscott et al., 2005; Yokoi et al., 2011). Our new framework predicts that tuning function changes driven by PD rotations have large systematic effects on subsequent sensorimotor learning.

A key idea is that PD rotation occurs not globally but rather only for neurons with PDs in a neighborhood of the movement direction associated with an experienced perturbation (Arce et al., 2010a, 2010b; Paz et al., 2003), because localized PD rotation will have anisotropic effects on both the spatial distribution of PDs and subsequent learning ability. In particular, localized PD rotation will confer a decreased PD density in directions that PDs rotated away from and an increased density in directions that PDs rotated toward. If neurons in the population displaying PD rotation act as learning units, a localized increase in PD density would then lead to greater recruitment of learning units for the associated movement direction, which should increase the rate of learning. Correspondingly, a localized decrease in PD density would lead to reduced recruitment of learning units for the associated movement direction, which should decrease the rate of learning. Therefore, our hypothesis makes the novel prediction that sensorimotor learning that elicits PD rotations inherently reshapes subsequent sensorimotor learning, as it would lead to systematically increased learning rates for movement directions near the destinations of PD rotations but systematically decreased learning rates for movement directions near the origins. This prediction is in stark contrast to conventional computational models where motor primitive tuning remains unchanged during learning, and no direction-dependent increases or decreases in learning unit density or learning rate are predicted (Donchin et al., 2003; Herzfeld et al., 2014; Hwang et al., 2003; Sing et al., 2009; Thoroughman and Shadmehr, 2000; Wainscott et al., 2005; Yokoi et al., 2011).

In this study, we develop a model of how the rotation of tuning curves induced by sensorimotor learning specifically influences subsequent sensorimotor learning ability, and in particular, the rate at which this subsequent learning can proceed. We test this adaptively-tuned learning model with behavioral experiments in humans and find a remarkable match to the precise biphasic pattern of increases and decreases in learning rates predicted by the properties of PD rotation. These results suggest that neural mechanisms underlying motor learning allow us to not only systematically alter the motor output we produce but also change the rate at which we alter this motor output (Braun et al., 2010; Harlow, 1949).

## RESULTS

### Predicting changes in the distribution of preferred directions driven by PD rotation

An accurate visuomotor map that transforms a visual target location in any given direction into a set of motor commands that can allow us to reach the target, is essential for reliable action generation (Hayashi et al., 2016; Wu and Smith, 2013) (Fig. 1a-c, light green line). For reaching arm movements, the PDs of neurons in M1, premotor cortex, and supplementary motor cortex have been shown to cover the entire range of movement directions (Fu et al., 1993; Lara et al., 2018; Scott et al., 1997; Scott and Kalaska, 1997), enabling the ability to generate reliable actions in any direction, as illustrated in Figure 1c (light green dots). For simplicity, we initially analyze the key predictions of PD shifts for motor learning ability using a uniform baseline distribution, and we later determine how these predictions change when the baseline distribution is anisotropic (Fig. S1).

**Figure 1:**
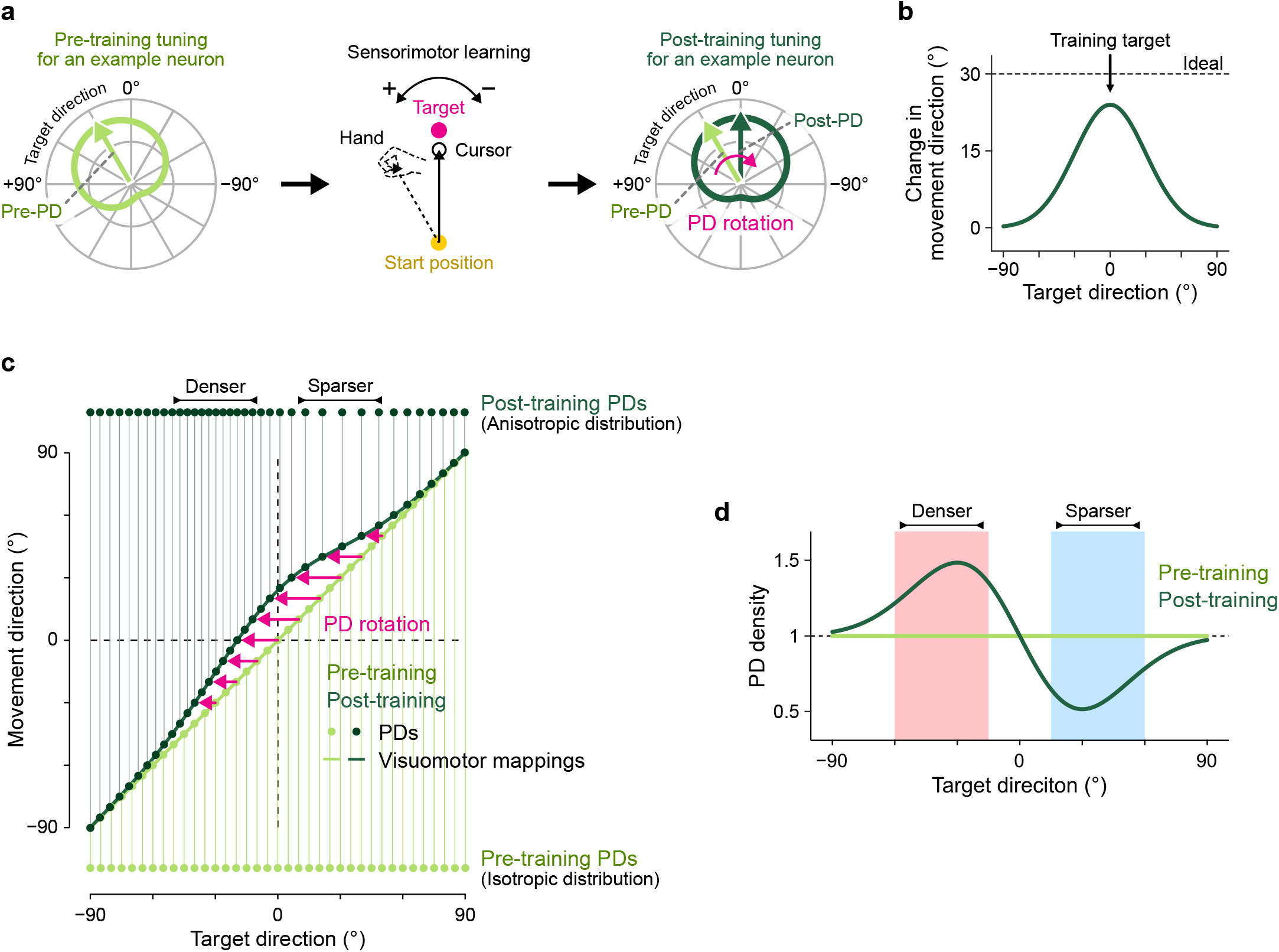
Visuomotor mapping predicts PD density. a. Schematic illustration of preferred direction (PD) rotation induced by sensorimotor learning. Neurons in motor cortical areas are tuned to the direction of movement, and the PDs of these neurons can rotate following sensorimotor learning. Consider a neuron with a baseline PD of +30° for target locations (left) that promotes hand movement in the +30° direction. To facilitate sensorimotor learning for which the +30° hand direction that the unit promotes becomes associated with a 0° target direction (center), this unit would tend to rotate its PD towards the 0° target direction (right). b. Generalization. The change in the hand movement direction conferred by the sensorimotor learning is local. It is maximal for the trained target and falls off for target direction farther from the training with a sigma of about 30° (Hayashi et al., 2016). c. Prediction of visuomotor mapping on PD rotation. At baseline, the hand movement direction would directly map to the target direction (light green line). However, in line with the local pattern of generalization, sensorimotor learning would locally deform the visuomotor map that relates movement direction to target direction (dark green line). Here, the changes in the target direction (pink arrows) associated with each movement direction would correspond to changes in PD, i.e. PD rotations, as illustrated. If the baseline PD were uniformly distributed (see the distribution of light green dots along the bottom), the local pattern of PD rotation would deform the PD density, resulting in an increased density at the destinations of PD rotations, and an increased sparsity at the origins of the PD rotations, as illustrated (see the distribution of dark green dots along the bottom). d. PD density. The non-uniform PD is predicted by the gradient of visuomotor mapping. The sensorimotor learning for the single target direction, which smoothly generalizes with a monophonic shape (b), changes the gradient from a unity (light green) to a biphasic shape (dark green).

Motor learning is known to distort the visuomotor mapping (Hayashi et al., 2016; Wu and Smith, 2013). Here we focus on the motor learning induced by visuomotor rotation (VMR), a directional perturbation where the cursor motion is skewed from hand motion (Fig. 1a). To correct the error, participants must learn to skew their hand motion in the opposite direction (Fig. 1a). Critically, adaptive changes from VMR training are locally generalized (Fig. 1b) and thus deform the shape of the visuomotor mapping rather than shifting it uniformly across target directions (Fig. 1c, dark green line). The specific distortion illustrated in Figure 1b is based on gaussian-shaped generalization with a sigma of 30° and 80% adaptation (Donchin et al., 2003; Hayashi et al., 2016; Krakauer et al., 2000). Neurophysiologic results indicate that the PDs of neurons in motor cortical areas rotate during motor learning (Alexander and Crutcher, 1990a, 1990b; Gandolfo et al., 2000; Li et al., 2001; Perich and Miller, 2017; Rokni et al., 2007; Shen and Alexander, 1997). Critically, this rotation corresponds to a horizontal shift in the visuomotor mapping for units local to the trained target direction (Fig. 1c, pink arrows and dark green dots). These localized horizontal shifts will change the density of the PD distribution, with an increase in density at the destinations of the PD rotations, in contrast to a decrease at the origins. Therefore, VMR training will transform an isotropic distribution of PDs at baseline into an anisotropic distribution following training, with a specific pattern of local increases and decreases from the baseline density.

Analytically, changes in PD density determine changes in the gradient (slope) of the visuomotor mapping, with a fixed shift across all PDs preserving a uniform distribution of PD density and leading to a global shift in the overall mapping that would maintain the unity slope of the baseline mapping. A generalization width of 30° would lead to a symmetrical 50% increase in the negative direction and 50% decrease in the positive direction of the baseline PD density, if the adaptation level is 80% for 30° VMR training as illustrated in Figure 1d. Critically, these changes in PD density would systematically alter the number of units activated for movements in different reach directions, with a greater number activated for reach directions with increased PD density and a smaller number for reach directions with decreased PD density. Therefore, if these units, or downstream consequences of their activity, were involved in subsequent motor learning, learning ability would be enhanced for target directions with increased PD density and reduced for target directions with decreased density. We would thus predict a pattern of learning rate changes that matches the biphasic pattern predicted for PD density changes.

### Modeling the effect of PD rotation on motor primitives and motor learning sensitivity

The task investigated in this study was a motor learning reaching task where a visual cursor representing the hand position was rotated around the starting position (VMR). A simple network model (Donchin et al., 2003; Herzfeld et al., 2014; Hwang et al., 2003; Sing et al., 2009; Thoroughman and Shadmehr, 2000; Wainscott et al., 2005; Yokoi et al., 2011) (Fig. 2a) can represent this learning based on an activity pattern (***g***) across directionally-tuned neural units, with the tuning of each unit centered at its PD (Fig. 2b, See Methods). Consider a single-trial motor adaptation paradigm with a fixed amount of visual error (*e*)

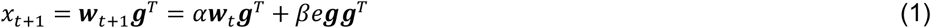

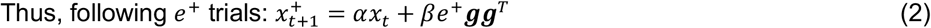

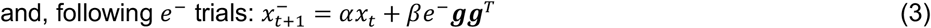

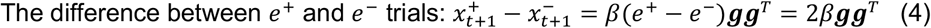

where the superscript *T* is the matrix transpose operator, the subscript *t* or *t* + 1 denotes the movement number, and *x* is the motor command (compensatory hand movement direction) to compensate for a VMR (*p*) and the motor error (*e*). Here *e*^+^ and *e*^−^ to denote positive and negative imposed errors, and show that the difference between the *x*_*t*+1_ values following *e*^+^ and *e*^−^ trials provides a measure of the learning sensitivity. The retention (*α*) and learning constants (*β*) determine the motor learning process. Equation 1 reveal that motor output in the next trial (*x*_*t*+1_) will depend on the amounts of motor memory with a retention constant (*αx*_*t*_) in addition to an error-dependent factor (*βe**gg***^*T*^) but Equation 4 combining the adaptive responses to *e*^+^ and *e*^−^ depends solely on the error-dependent factor, which can ignore the amounts of motor memory (*x*_*t*_) across target directions.

**Figure 2:**
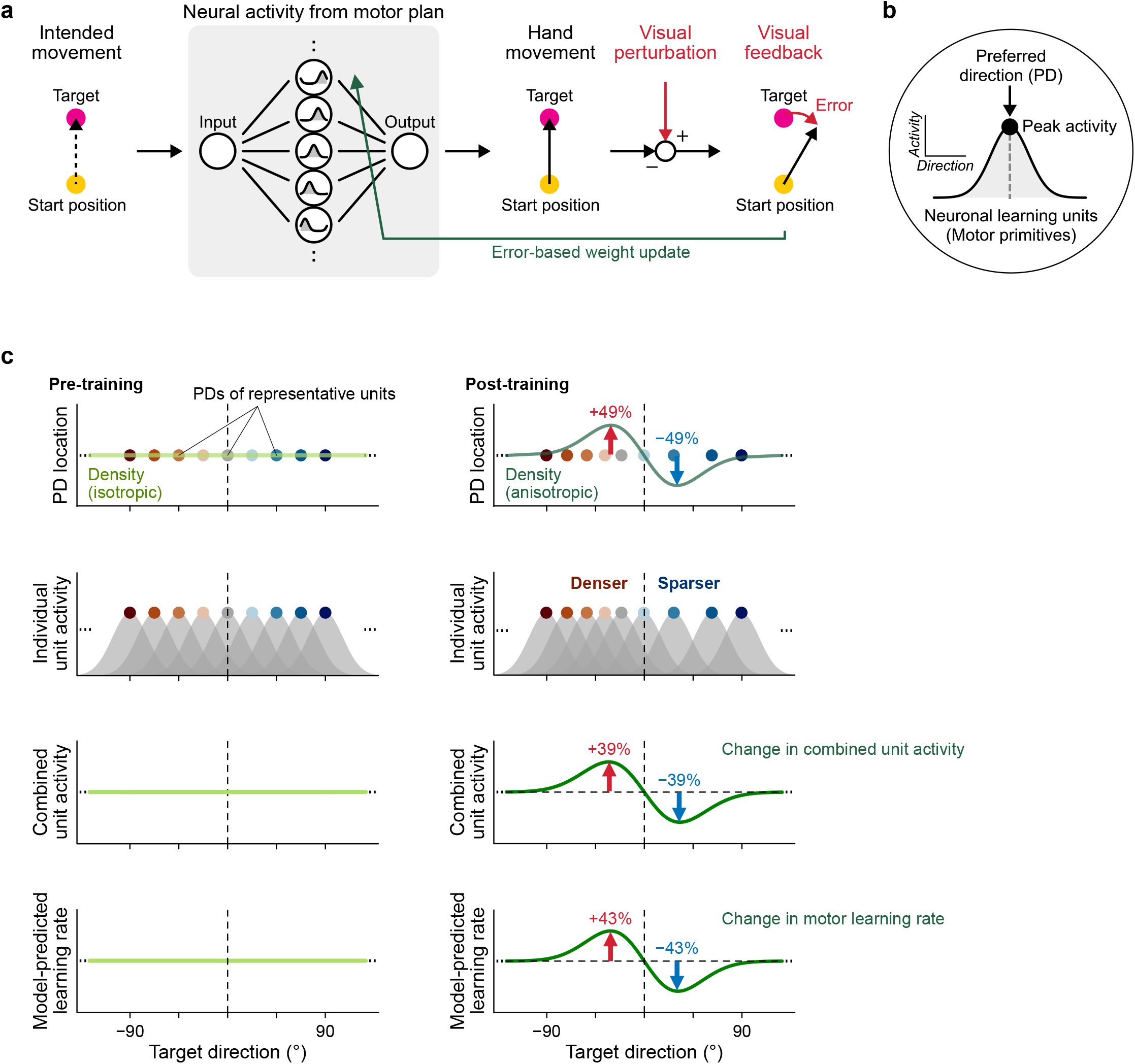
The adaptively-tuned learning model predicts the effects of PD rotation on the motor learning rate. a-b. Modeling the effect of PD rotation on sensorimotor learning. In this model neural units display activity patterns that show preferred direction (PD) tuning, and these units act as motor primitives in that their outputs are combined to form motor commands. Sensorimotor perturbations result in motor errors that induce weight updates for how the output of these units are combined that act to reduce experienced errors. c. Modeling results for the effect of sensorimotor learning on the spatial patterns of PD density, neural activity, and learning rate. 1st row: Sensorimotor learning induces local PD rotations that change the uniform distribution of PD density present at baseline to a biphasically-shaped pattern as also illustrated in Figure 1c. This biphasic pattern arises from decreases in PD density for PDs at the origins of the PD rotation and increases in PD density for PDs at the destinations of the PD rotation. 2nd & 3rd rows: The learning-induced changes in PD density result in corresponding changes in neural activity elicited by each target direction. 4th row: The spatial changes in neural activity result in corresponding changes in motor learning rate.

We next used the pattern of PD rotations elicited by training a 30° VMR (Fig. 1b). A diagram of the effect of this training is illustrated in Figure 2a, showing that perturbations arising from VMR training lead to visual feedback of error that elicits motor plasticity. In particular, as diagrammed in Figure 1c and d, PD rotations induced by VMR learning result in increases in PD density in the target directions corresponding to the destinations of PD rotations and decreases in PD density in the target directions corresponding to their origins. The biphasic anisotropy of this post-training PD density would then predict a similarly anisotropic but slightly broader biphasic pattern of neural activity, smoothed by the finite-width tuning curves of these cells. If these same neural units were responsible for subsequent motor learning, the anisotropic pattern of neural activity across movement direction would, in turn, predict a specific pattern of changes in the sensitivity of motor learning corresponding to the ***gg***^*T*^ squared neural activity dependence in Equation 4. All three consequences of a learned visuomotor adaptation that arise from our model – the biphasic patterns of PD density, neural activity, and critically, motor learning sensitivity are plotted out in Figure 2c. These patterns are only subtly different from one another as they all arise from the same direction-specific pattern of PD rotation that results from local visuomotor learning (Fig. 1b), and therefore, all display the same striking biphasic appearance.

Another notable prediction of this adaptively-tuned learning model is that the average motor learning rate across all target directions would be unchanged from baseline following PD rotation, as the predicted increases and decreases would be balanced. This is in line with preservation of the units that contribute to motor learning in our model. The adaptively-tuned learning model, as neither creates nor abandons motor primitives but rather reorganizes the PD distribution and thus refocused these units based on local PD rotations.

Note that the simulation results shown in Figure 2, are based on a uniform distribution of PD’s at baseline, but the specifics of the predicted change in the PD distribution would be influenced by the baseline PD distribution. Based on neurophysiologic data during primate reaching movements, there is believed to be an anisotropy in the baseline distribution of PDs that displays an elliptical shape, where the long axis is closely aligned with the orientation of the forearm (Scott et al., 2001). The long axis of this ellipse corresponds to the axis in which the inertia of the arm is greatest, suggesting that PD density approximately matches the force required to move the arm in each direction (Gribble and Scott, 2002; Herter et al., 2007; Scott et al., 2001, 1997; Scott and Kalaska, 1997). We find, however, that this elliptical pattern for the baseline PD distribution has only subtle effects on model-predicted changes in PD density for a wide range of elliptical anisotropy levels (Fig. S1).

### Experimental results show large changes in motor learning sensitivity across target directions

We designed an experiment in humans to test the prediction that the pattern of PD rotation that would arise from the adaptation to a sensorimotor perturbation would lead to a large biphasic pattern of increases and decreases in the rate of subsequent motor learning. Following a baseline period where participants became familiar with the basic point-to-point reaching arm movement task, they adapted their movements toward a frontal target (0°) when a sensorimotor perturbation was imposed (Fig. 3a and b). This perturbation was generated by a VMR that was gradually introduced by being slowly ramped up in amplitude from 0° to 30° (0.25°/trial) to reduce the use of explicit strategy compared to an abrupt-onset perturbation (Modchalingam et al., 2023). This rotation was applied in either the clockwise (CW) or the counter-clockwise (CCW) direction – counterbalanced across participants (Fig. 3c, CW group: N = 11, final rotation of −30°; CCW group: N = 11, final rotation of +30°). Data from participants who learned CW and CCW VMRs were combined by flipping data from the CW group (see Methods). As expected, adaptation to the VMR perturbation induced adaptive changes in the hand movement direction (i.e., the learning direction) opposite to the direction of VMR (i.e., the intervention direction) that compensated it. Critically, the participants performed test-blocks with pairs of perturbation and probe trials (Fig. 3d), both before and after this training, in order to measure changes in the sensitivity of motor learning to these single-trial perturbations. On the perturbation trials, participants performed reaching movements toward one of the test targets (0°, ±15°, ±30°, ±60°, ±90°, and 180°, Fig. 3b) with a cursor clamp that constrained the cursor motion to be skewed from the target direction by 7.5° in either the CCW or CW direction (i.e. ±7.5°), randomly assigned independently of the hand movement direction (Morehead et al., 2017; Vaswani and Shadmehr, 2013) (Fig. 3d). In the subsequent probe trial, the participants performed a reaching movement toward the previous target with zero-error cursor clamped feedback (Fig. 3e), so that the cursor motion was always directed straight to the target. This allowed us to measure any systematic perturbation-related changes in movement direction in the participant’s hand motion between the perturbation trial and the ensuing probe trial to assess perturbation-driven learning (with compensatory changes, opposite in direction to the visual perturbation, defined as positive).

**Figure 3:**
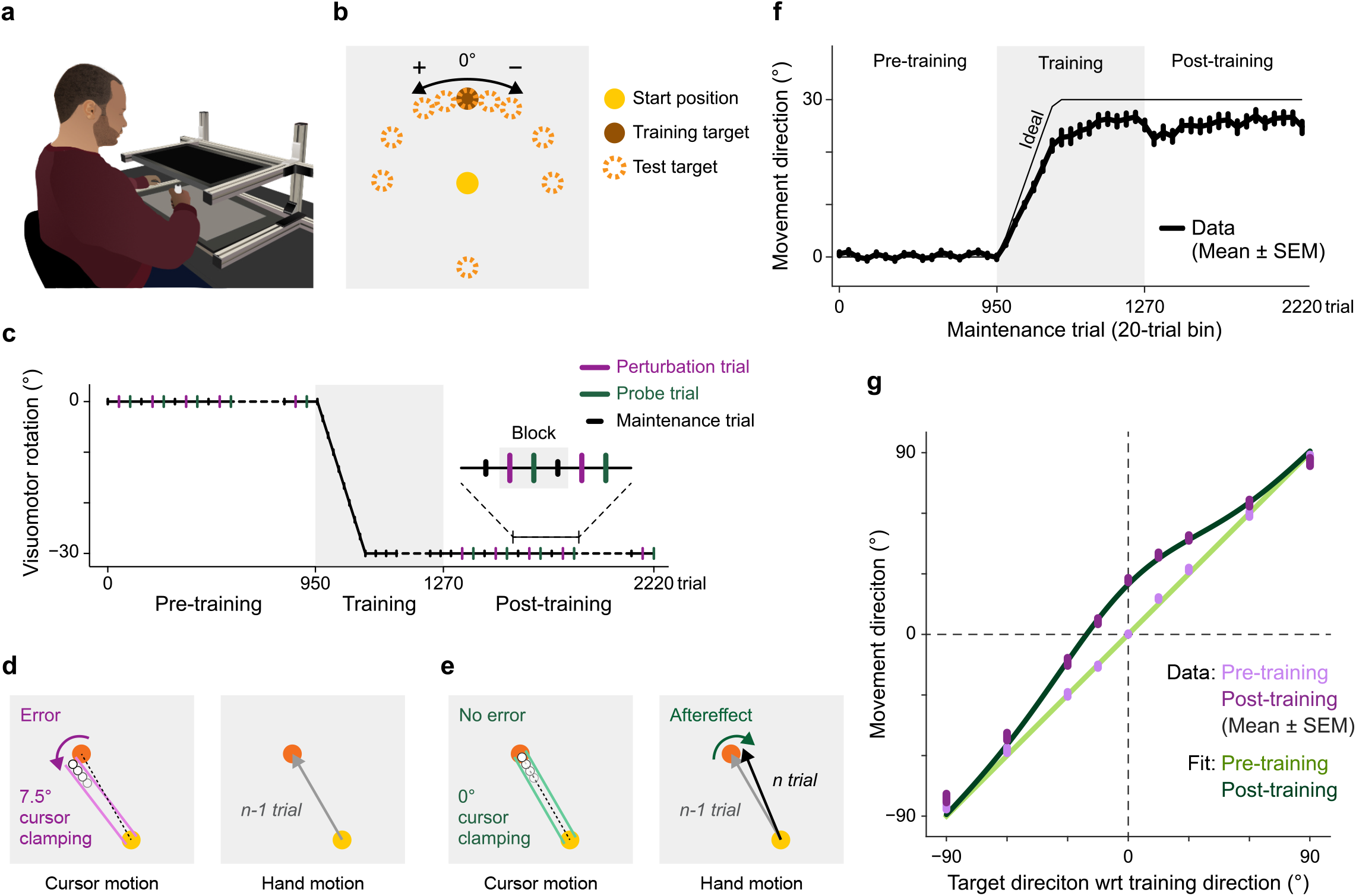
Experiment diagram and learning-induced changes in the visuomotor map. a. Experiment diagram. Participants performed visually-guided reaching arm movements from a start circle to one of several possible target locations when cued by target appearance. b. Target locations. Sensorimotor learning was trained only at the frontal target location (pink circle). The green dotted circles indicate the 10 possible probe target locations used to assess how motor learning rate varied across target directions in the pre- and post-training periods. c. Trial sequences. Participants adapted to either a CW or CCW visuomotor rotation (VMR) while reaching toward the training target. The VMR was gradually increased over the span of 120 trials from 0° to -30° (CW) or +30° (CCW) with the CW rotation illustrated here. Before and after training, a series of test pairs, each consisting of a perturbation trial followed by a probe trial in one of the 10 probe directions were randomly interleaved with one another. d-e. Measurement of motor learning sensitivity. Participants iteratively performed a series of paired (d) perturbation and (e) probe trials at each probe location. In the perturbation trial (d), they were exposed to a CW or CCW 7.5° error clamp perturbation (chosen at random) at the probe target (pink), with the CW perturbation illustrated. This 7.5° error clamp perturbation imposed a fixed-amplitude 7.5° CW or CCW visual error (red arrow). In the subsequent probe trial (e), participants again performed a reaching movement to the same probe target, but with a zero-error cursor clamp. This allowed us to measure any changes in post-perturbation movements (green arrow). f. The learning curve for the gradual onset ±30° VMR training. Each data point shows the population-averaged movement direction for a 20-trial bin for movements to the trained frontal target location. g. Visuomotor mapping. The hand movement direction for the perturbation trial shows how the effect of the sensorimotor learning associated with the ±30° VMR training intervention was generalized to reach movements to the neighboring targets. The relationship between the target and the hand movements (light and dark purple for the pre- and post-training phases, respectively) was similar to that predicted in Figure 1a. The light and dark green lines show Gaussian fits to the observed generalization patterns.

We found that the gradual 30° VMR training resulted in a 25.16° ± 0.93° adaptation (Fig. 3f) that generalized narrowly to the movements for neighboring targets (purple lines in Fig. 3g), corresponding to a locally-deformed the visuomotor map as predicted (green lines in Fig. 3g). Notably, we found no shift on the center of the deformed visuomotor mapping that would arise from explicit aiming (Fig. S2) (Day et al., 2016), suggesting the VMR training paradigm we used primarily resulted in implicit learning, with little use of explicit strategy. When examining the test-block data before VMR training, we found motor learning rates to be fairly consistent across movement directions and highly symmetric around the 0° training direction for the subsequent VMR training (Fig. 4a – 3.73° ± 0.25°, mean ± SE across directions and also across participants for the mean across directions). In contrast, motor learning rates became markedly asymmetric around the 0° training direction following the VMR perturbation training. Learning rates increased in the perturbation direction but decreased in the learning direction as predicted by our model of learning-driven changes in the PD distributions for learning units (Figs. 2c and 4a). Notably, the overall motor learning rate averaging across all probe target directions remain unchanged (Fig. 4b, 3.73° ± 0.25° vs 3.91° ± 0.23°, t(21) = 1.22, p = 0.24) before vs after the VMR training. but the difference of motor learning rate between the increased (red) and decreased (blue) sides (Asymmetry index) was found not in pre-training phase (−0.01 ± 0.10, t(21) = 0.13, p = 0.90) but in the post-training phase (0.39 ± 0.07, t(21) = 5.56, p = 1.63 × 10^−5^), and statistically increased it from pre-to post-training phases (Fig. 4c, t(21) = 4.78, p = 1.02 × 10^−4^), which is in line with our prediction from the adaptively-tuned learning model (Fig. 2). Specifically, we found that the post-training – pre-training difference (Fig. 4d) was remarkably similar to both the shape (r = +0.85 for isotropic prediction, r = +0.89 for anisotropic prediction) and amplitude (peak-to-peak modulation, 75.6 ± 16.2% for experimental data, 77.8% for the isotropic prediction, and 71.3% for the anisotropic prediction) of the theoretical prediction from Figure 2c. Moreover, when we regressed the experimental data onto the biphasic model-predicted pattern of changes in learning rate, we found regression coefficients that were near unity and significantly above zero for models based on both isotropic and anisotropic baseline PD distributions (regression coefficient = 0.78 ± 0.13, t(21) = 5.77, p = 9.98 × 10^−6^ for isotropic prediction, 0.95 ± 0.14, t(21) = 6.69, p = 1.27 × 10^−6^ for anisotropic prediction) (Fig. 4e). In line with the predicted pattern, the data showed both increased learning rates in the perturbation direction (33.0 ± 6.4%, t(21) = 5.17, p = 3.97 × 10^−5^), decreases learning rates in the learning direction (−15.7 ± 5.9%, t(21) = 2.56, p = 0.0184) and significantly positive differences (+48.7 ± 8.0%, t(21) = 6.05, p = 5.25 × 10^−6^) (Fig. 4f). Together, these findings provide compelling evidence for the prediction that PD rotation of learning units determine subsequent motor learning rates (Figs. 1 and 2).

**Figure 4:**
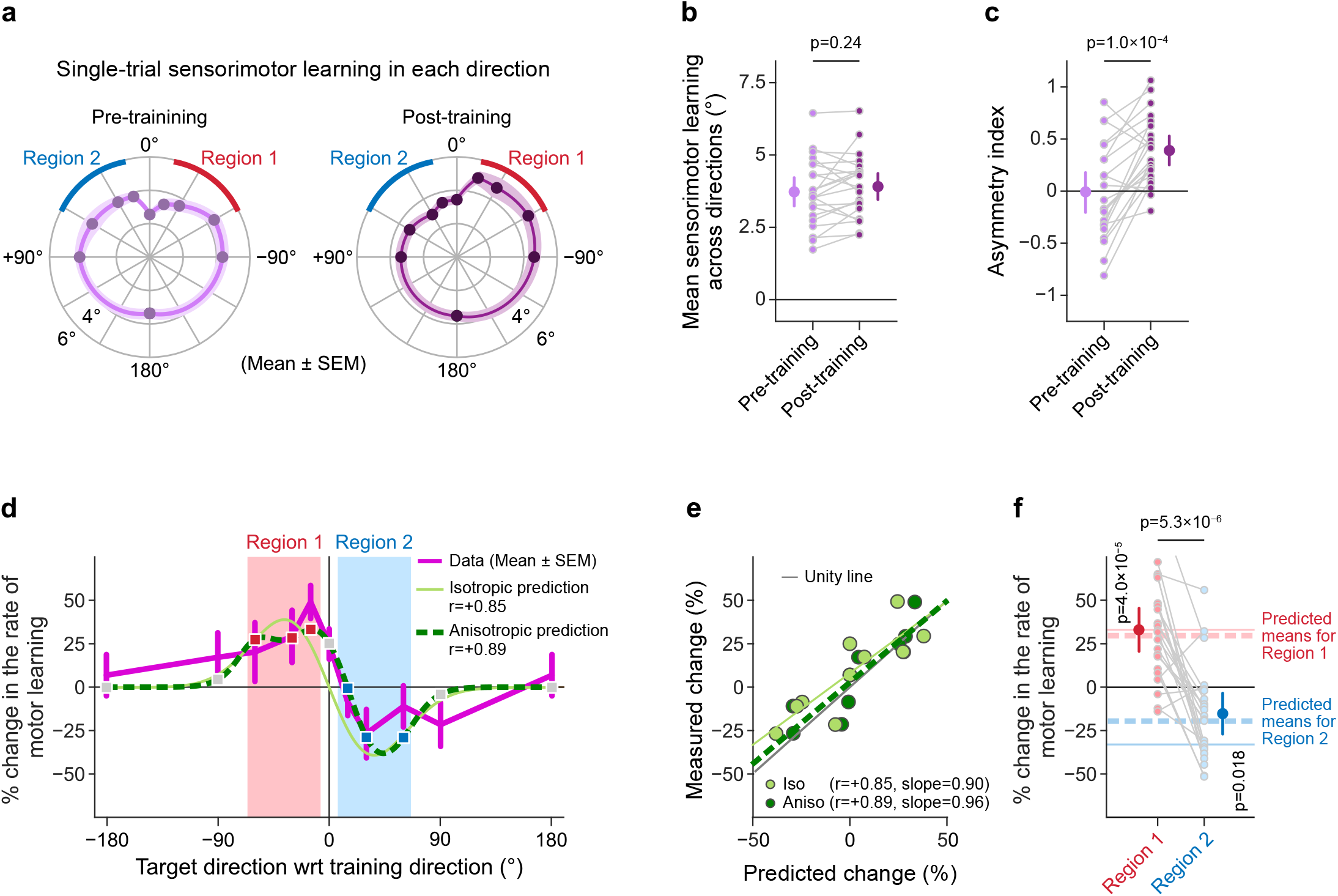
Influence of motor learning intervention on sensorimotor learning rates. a. Sensorimotor learning rates before and after VMR training. Polar plots indicate how the motor learning sensitivity depends on the target directions to be reached. The motor learning rate was largely symmetric around the frontal target before the ±30° VMR training intervention, but after the intervention it became markedly asymmetric, as it increased in the perturbation (region 1 colored with red) and decreased in the learning direction (region 2 colored with blue). b. Preservation of direction-averaged motor learning sensitivity. When averaged across probe targets, motor learning sensitivity was not different pre-vs post-training. Each data point indicates the data from a single participant. c. Systematic changes in the symmetry of motor learning sensitivity. The plotted asymmetry index indicates the difference in motor learning sensitivity between the red and blue regions illustrated in panel (a). This asymmetry index was not different from zero in the pre-training phase, but it became > zero in the post-training phase, as predicted. d. Changes in the spatial pattern of motor learning sensitivity. Sensorimotor learning rates increase for some target directions and decrease for others from pre-to post-training. The spatial pattern of these changes (purple) closely matches the bi-phasic patterns predicted by both versions of the adaptively-tuned learning model, with an isotropic baseline (light green) and with an anisotropic baseline corresponding the Scott et al data (Scott et al., 2001) (dark green). Note that these versions make only subtly different predictions, and are both highly correlated with the data. e. Comparison of predicted and measured changes in motor learning for isotropic-baseline (light green) and anisotropic-baseline (dark green) versions of the model. Measured and predicted changes are closely matched, lying near the unity line for both versions of the model (r=0.85-0.89, slopes=0.90-0.96). f. Summary of the effects of PD rotation for subsequent sensorimotor learning. The means of the learning rate changes in region1 and region 2 are higher and lower than zero, respectively, in line with the bi-phasic pattern predicted by the adaptively-tuned learning model. Error bars in (a and d) show SEM, and in (b, c, and f) show 95%CI.

## DISCUSSION

PD rotation has been broadly reported for neurons in motor cortical areas during motor learning (Alexander and Crutcher, 1990a, 1990b; Gandolfo et al., 2000; Li et al., 2001; Perich and Miller, 2017; Rokni et al., 2007; Shen and Alexander, 1997). In these studies, monkeys adapted reaching movements in novel kinematic (e.g., visuomotor rotation) or dynamic (e.g., force-field) environments. Almost all studies have considered that PD rotation reflects changes in movement parameters (i.e., the outcome of motor learning). For example, studies have considered that PD rotation is associated with altered electromyography (Kadota et al., 2014; Li et al., 2001) or dynamics (Rokni et al., 2007) of the arm being controlled. However, a recent study showed that PD rotation does not fully account for these behavioral changes, as motor learning continues even after PD rotations are complete, suggesting that adaptive changes downstream of PD rotations may play a key role in sensorimotor learning (Perich and Miller, 2017).

Here, we suggest that the spatially-anisotropic changes in the PD distribution resulting from local PD rotation can have a dramatic spatially-specific effect on learning ability (Figs. 1 and 2). The model for sensorimotor learning that we propose differs from conventional ones, in that they assume that spatially-invariant learning units provide the substrate for sensorimotor learning (Donchin et al., 2003; Herzfeld et al., 2014; Hwang et al., 2003; Sing et al., 2009; Thoroughman and Shadmehr, 2000; Wainscott et al., 2005; Yokoi et al., 2011). In contrast, our new model, is based on the idea that learning units that sculpt adaptive changes in motor output are retuned with the PD rotation that occurs during motor learning. This PD rotation alters the spatial distribution of preferred directions by producing increases in PD density at the destinations of the rotation and decreases at the origins, with these density changes dictating changes in the numbers of learning units that are recruited for movements in different directions. Thus, broadly, our adaptively-tuned learning model dictates that sensorimotor learning changes not only the motor command that is produced but also the way in which it is learned (Figs. 1 and 2).

Specifically, our adaptively-tuned learning model predicted a striking biphasic pattern of changes in learning rates across different movement directions following sensorimotor learning (Fig. 2c). We, therefore, designed an experiment to determine whether the strength of adaptive responses would indeed be systematically altered by sensorimotor learning as predicted (Fig. 3). The results show not only that sensorimotor learning induces large systematic changes in learning rates (Fig. 4), but also that the spatial pattern of these changes closely matches (r ≥ +0.85) the biphasic pattern predicted by our adaptively-tuned learning model (Fig. 4d).

### Computational implications

Our hypothesis, that changes in PD density determine the rate at which motor learning proceeds, can be viewed computationally as dictating a direct link between the rate at which sensorimotor learning occurs for each target direction and the gradient of the visuomotor map at that same target direction. This is because changes in the gradient of the visuomotor map that defines how motor output depends on visual information about target location (Fig. 1c) directly correspond to local changes in PD density (Fig. 1d). Therefore, the rate of sensorimotor learning directly relates to the amplitude of the mathematical derivative of the motor output with respect to visual information about target direction. However, this derivative is precisely the inverse of the derivative of target error with respect to motor output, i.e. the error derivative (Abdelghani et al., 2008), that dictates gradient descent learning for optimizing motor output to minimize performance errors. This makes the PD density driven changes in the sensorimotor learning rate that our hypothesis dictates, inversely proportional to the step sizes dictated by the popular gradient descent learning rule (Rumelhart et al., 1986) that is widely used in modern machine learning applications (LeCun et al., 2015).

The disparity between the learning rate changes that we observe and those predicted by gradient descent learning is surprising, and might suggest that both the changes in sensorimotor learning that we uncover and the theory that predicts them result in learning that is, computationally, highly suboptimal. However, the inverse gradient learning rate that our hypothesis predicts precisely corresponds to the step size that would be dictated by Newton’s method (also referred to as Newton-Raphson) for finding zero-error solutions (Ypma, 1995). A key difference between Newton’s method and common gradient descent, where the step size is directly rather than inversely proportional to the gradient of error with respect to motor output, is that Newton’s method is superior when in the neighborhood of an existing zero-error solution. The crux of Newton’s method is that for a given error level in this neighborhood, a steeper gradient between the amount of error reduction and the step size would dictate that a proportionately smaller step would be needed to arrive at the zero-error solution, whereas proportionately increasing the step size with the gradient, as called for by common gradient descent, would make overshooting this solution more likely. This is an issue for sensorimotor learning, which often operates in the neighborhood of zero-error solutions. However, it is not an issue, when far away from a zero-error solution, where complex high-dimensional machine learning problems operate. In fact, modern deep learning models are often intentionally stopped long before they can converge on a zero-error solution for the training data, because this early stopping has been shown to reducing overfitting and thus improve generalization. Thus, the mechanism we uncover shows not only that changes in neural tuning can have profound effects on sensorimotor learning ability but also that, computationally, the observed changes in sensorimotor learning reflect the difference in how learning rules should operate when in the neighborhood of a zero-error solution vs far from it.

### Progress in understanding mechanisms for controlling sensorimotor learning rates

The ability to learn is perhaps the defining feature of the nervous system, and plethora of studies have focused on documenting changes in neural activity that result from learning (Gandolfo et al., 2000; Li et al., 2001; Medina and Lisberger, 2008; Paz et al., 2003; Suvrathan et al., 2016). However, some recent studies have made more fundamental advances into understanding the neural basis of sensorimotor learning itself. For example, multielectrode recordings in monkeys are beginning to elucidate the relationships between planning-related activity in premotor areas and learning ability (Perich et al., 2018; Sun et al., 2022; Vyas et al., 2018).

Still the mechanisms that control that rate at which learning-related changes in neural activity and motor output occur remain poorly understood. However, that is just beginning to change. Using two-photon calcium imaging and optogenetic control of activity in Drosophila, dopamine release associated with turning behaviors has been shown to increase the rate of error-dependent sensorimotor learning (Fisher et al., 2022; Grover et al., 2022), and the current study demonstrates that sensorimotor learning alters the PD distribution that in turn determines the rate of subsequent learning. Perhaps, the tide is now turning and we are on our way to more wholly understanding this most elegant of abilities, the ability to shape learning by controlling the rate at which learning occurs.

## METHODS

### Participants

22 participants (aged 20-31 years, 10 male and 12 female) with no known neurological impairment were recruited to perform the following behavioral experiments. Written informed consent was obtained from all participants prior to the experimental sessions, and the experiments were approved by the Harvard University committee on the use of human subjects.

### Experimental procedures

The participants used their right hands to hold a 2.5cm-diameter handle that encapsulated a digitizing pen and slid it on along the surface of a Wacom tablet (the sampling rate was 200 hz) that recorded hand motion (Fig. 3a). They were seated in front of a horizontal display over the tablet that prevented them from directly viewing their own arm.

At the beginning of each trial, a red target (diameter: 1 cm) appeared 10 cm from the start circle (diameter: 1 cm). The participants were required to move a green cursor (diameter: 3 mm) representing the hand position toward the target as straight as possible. A warning sound about the long movement duration dinged at the completion of each trial if it was above 500 ms. After the completion, the green cursor was replaced with the green open circle whose radius was matched with the distance from start position to guide a hand position towards the start position without visual errors. Participants were asked to always aim at the target directions and not to use their explicit strategy.

Prior to the main experiment, the participants performed practice trials for 0°, ±30° ±60°, ±90°, ±120°, ±150°, and 180° targets to familiarize themselves with the reaching movement task (Fig. 3b). The main experiment then consisted of three phases: a pre-training phase, a training phase, and a post-training phase (Fig. 3c). In the training phase, half of the participants adapted to a 30° counterclockwise (+30°) visuomotor rotation (VMR) and the other half to a 30° clockwise (−30°) VMR. The VMR training was provided for the 0° target away from the body in the midsagittal plane (Fig. 3b). This training began with a gradual 0.25° per trial increase in the VMR, from 0° to 30° (121 trials). This ramp-up was followed by 195 “extended intervention” trials with a 30° VMR (195 trials, Fig. 3c) to maintain asymptotic adaptation.

In the pre-training and post-training phases, the participants iteratively performed short three-trial blocks. Each consisted of a pair of perturbation and probe trials (Fig. 3d and e) to one of the test targets (0°, ±15°, ±30°, ±60°, ±90°, and 180°, Fig. 3b) followed by a training trial with null VMR in the pre-training phase and with 30° VMR in the post-training phase to the training target (0°). In the perturbation trial, a cursor clamp perturbation (Morehead et al., 2017; Vaswani and Shadmehr, 2013) was applied, in which the cursor’s direction was deviated by ±7.5° from the test target, randomly ordered. This enabled us to impose identically-sized visual errors before and after the sensorimotor learning in any target direction. In the subsequent probe trial, the participants performed reaching movements toward the same test target, but with a zero-error cursor clamp (i.e., 0°) in order to measure the aftereffect (motor learning rate, see below section) induced by the ±7.5° cursor clamp perturbation. In total, participants performed 30 perturbation-trial / probe-trial pairs (15 pairs with +7.5° and 15 pairs with −7.5° perturbations) for each of the 10 test targets in both the pre- and post-training phases.

The 320-trial training phase was administered in 2 blocks, and 950-trial the pre- and post-training phases were administered in 10 blocks each, with blocks separated by rest breaks that were typically 1-2 minutes in duration. Each block in the pre- and post-training phases began with 5 retraining trials to the training target to refamiliarize participants with the current perturbation level.

## Data analysis

We measured the hand movement direction at the peak speed point of each movement in order to investigate the generalization, visuomotor mapping, and motor learning rate. The sign of this movement-direction data was reversed for training trials with CW sensorimotor learning to align the CW and CCW data with respect to the trained VMR. We used a threshold of 0.05 for significance in all statistical tests.

### Sensorimotor learning

We measured the amounts of the sensorimotor learning. We gathered the data with every 20-trial bin of the intervention trials and computed the mean after an outlier removal within each bin (Fig. 3f). The criterion of the outlier removal is 3IQRs away from the median, which is almost 0.005% chance if it is sampled by normal distribution.

### Generalization and visuomotor mapping

We computed the mean of the hand movement directions in the perturbation trials for each of the test target directions in each of the pre- and post-training phases after removing outliers that is 3IQRs away from the median (Fig. 3g).

We assumed that baseline visuomotor mapping is sufficiently accurate, i.e., *x*(Δθ) = Δθ, then modeled the acquired visuomotor mapping (*x*) with a gaussian function (*G*) of movement direction with respect to the training direction (Δ*θ*) as follows: *x*(Δθ) = Δθ + G(Δθ) = Δ*θ* + λexp{ − Δ*θ*^2^/2σ^2^}, *λ* is the amount of learning and *σ* is the generalization width. We fitted the model into the acquired visuomotor mapping to get them. The fitted visuomotor mapping implies how the PD of learning units was shifted. Because the PD is thought to remain constant in the hand movement direction, the PD is shifted at the target direction after the visuomotor mapping is distorted (Fig. 1c). Theoretically, the changes in the PD density are predicted by the gradient of visuomotor mapping (Fig. 2f).

Furthermore, we also utilized the generalization curve to investigate how much our behavioral results were contributed by explicit strategy. Previous studies showed that the center of generalization is not directed towards actual target directions but intended movement direction (Day et al., 2016), suggesting that the degrees to shifting the center is a measure of the explicit strategy. We fit a gaussian function with an additional parameter, the center shift (*θ*_*cs*_), for each individual generalization curve as follows: *G*_*cs*_(Δθ) = λexp{−(*θ*_*cs*_ − Δ*θ*)^2^/2*σ*^2^}. To test that the center of generalization was biased towards the intended movement direction, we performed a one-sample t-test against 0° target directions (Fig. S2).

### Motor learning rate

We investigated how sensorimotor learning influenced the subsequent motor learning rate by comparing predictions of our adaptively-tuned learning model to experimental data for learning rate changes (Fig. 2). We computed the sensorimotor learning rate in our experimental data by subtracting the hand movement direction in the perturbation trial from that in the probe trial (Δ*x*(*θ*)) and then normalizing this difference by the perturbation experienced, as follows: −changes in the motor learning rate 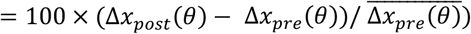

### Computational model

We adopted one of the prevailing models for sensorimotor learning: the motor primitive model (Donchin et al., 2003; Herzfeld et al., 2014; Hwang et al., 2003; Sing et al., 2009; Thoroughman and Shadmehr, 2000; Wainscott et al., 2005; Yokoi et al., 2011) (Fig. 2a). This model accounts for motor learning behaviors with learning units (***g***) that are recruited according to the tuning function of the target direction (*θ*), typically expressed by a gaussian radial basis function: *g*_*i*_(*θ*) =exp {−(*θ* − *φ*_*i*_)^2^/2σ^2^} (***φ*** = [*φ*_1_, *φ*_2_, …, *φ*_*N*_] is the PD and *σ* is the gaussian tuning width) (Donchin et al., 2003; Herzfeld et al., 2014; Hwang et al., 2003; Sing et al., 2009; Thoroughman and Shadmehr, 2000; Wainscott et al., 2005; Yokoi et al., 2011) (Fig. 2b). The motor learning behaviors occur (Fig. 2a) as follows:

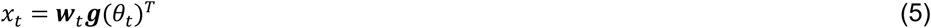

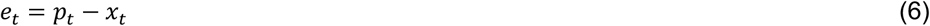

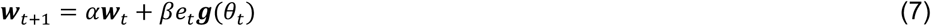

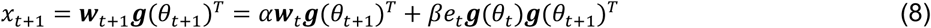

The motor command (*x*) is determined by the activity of the motor primitives (***g***(*θ*) = [*g*_1_(*θ*), *g*_2_(*θ*), …, *g*_/_(*θ*)]) and the weighting parameter (***w*** = [*w*_1_, *w*_2_, …, *w*_*N*_]) (*N* is the number of motor primitives) (Eq. 5). When a movement error (*e*) is produced by exposure of the VMR (*p*) (Eq. 6), the motor learning system updates the weights (***w***) according to the movement error (Eqs. 7 and 8). *α* and *β* are retention (close to but less than unity) and learning constants (*β* > 0), respectively.

### Model prediction testing and statistics

We predicted modulation of the motor learning rate with the motor primitive framework (Donchin et al., 2003; Herzfeld et al., 2014; Hwang et al., 2003; Sing et al., 2009; Thoroughman and Shadmehr, 2000; Wainscott et al., 2005; Yokoi et al., 2011) (Fig. 2a). Because the weight updates in the model are dependent both on motor errors (*e*) and the activity of the neural units (***g***) (Eq. 7), the rate of error-dependent sensorimotor learning depends on the squared sum of the neuronal unit activities (Eq. 8). Thus, the PD rotation, that differently regulates the unit activities across target directions, should influence the motor learning rates. Here, we incorporated it into the model and demonstrated whether the model prediction with PD rotation is consistent with the behavioral results.

The model with PD rotation is based on the neurophysiological findings. The PDs are assumed to be uniformly distributed across all target directions at baseline (Fu et al., 1993; Gribble and Scott, 2002; Herter et al., 2007; Lara et al., 2018; Scott et al., 2001, 1997; Scott and Kalaska, 1997) and are rotated with sensorimotor learning (Alexander and Crutcher, 1990a, 1990b; Gandolfo et al., 2000; Li et al., 2001; Perich and Miller, 2017; Rokni et al., 2007; Shen and Alexander, 1997). The PD rotation is believed to reflect the changes in motor outputs, i.e., the resultant PD would change into a target direction where the same movement direction is accomplished. Thus, the PD rotation will locally shift at the horizontal axis on visuomotor mapping diagram (Fig. 1c), which leads to biphasic changes in PD density. An example generalization/visuomotor mapping shown in Figure 1 will increase and decrease 49% in the PD density (1st row). This modulates individual unit activities (2nd row), and then increases and decreases the sum of unit activities 39% (3rd row in Fig. 2c). The motor learning rate, that is, squared sum of unit activities increases and decreases 43% (4th row in Fig. 2c). Importantly, their biphasic shapes are mostly identical although the peak and its location is slightly different. Note that some studies showed that PD density at baseline is skewed (Gribble and Scott, 2002; Herter et al., 2007; Scott et al., 2001, 1997; Scott and Kalaska, 1997), but we discussed that there are relatively very small effects on the following model prediction in supplementary materials (Fig. S1)

There are three key predictions to be tested here. First, since the changes in PD density that drive learning rate changes are entirely local in that the total number of learning units is unchanged, the model predicts that overall sensorimotor learning rates averaged across all target directions should remain constant. To examine this prediction, we computed average motor learning rates across all target directions both pre- and post-training and compared them using a paired t-test. Second, the model predicts a biphasic spatial pattern of learning rate modulation whereby learning rates increase at the destinations of PD rotations but decrease at the origins of these rotations. We defined positive and negative areas as [−15°, −30°, −60°] target directions and [+15°, +30°, +60°] target directions, respectively, and examined the asymmetry by computing ([positive areas] – [negative areas]) / [mean of them] (asymmetric index) with one-sample t-test. Third, the biphasic modulation of the motor learning rates is predicted by PD density. To do so, we first estimated PD density with the curve fitting into the mean visuomotor mapping (see Generalization and visuomotor mapping section). Then, we introduced the novel PD density into the motor primitive model and predicted motor learning rate. This change in the motor learning rate thus reflected the actual visuomotor mapping in our dataset. We computed correlation coefficients between predicted and actual patterns of % changes in motor learning rate as well as numerically compared them at the positive and negative areas with one-sample t-test. Furthermore, to examine the individual differences, we performed the aforementioned process for each individual data, and performed linear regression from predicted to measured changes.

## Acknowledgements

We gratefully thank the members of the Nozaki, the Takiyama, and the Smith laboratories for their in-depth and insightful comments. We are also grateful to Asako Munakata, Yasuko Shinya, Kanae Abe, and Sarah Burrous for managing our behavioral experiments. This study was supported by the following grants: the Uehara Memorial Foundation Postdoctoral Fellowship, JSPS Postdoctoral Fellowship for Research Abroad (JP202160367) and JSPS KAKENHI (JP23H03296) and JST/PRESTO (JPMJPR23S8) to TH, National Institute of Neurological Disorders and Stroke R01 (NS105839) and Nation Science Foundation (NSF 2218427) to MAS, and JSPS KAKENHI (JP17H00874, JP18H04082, JP18H03143, JP18H03154, JP20H05459, JP21H04860, JP22K19736) to D.N.

## Data availability

The data in the current study are available upon reasonable request.

## Code availability

All simulation and analysis code are available upon reasonable request.

## Supplementary materials

**Supplementary Data Figure 1.**
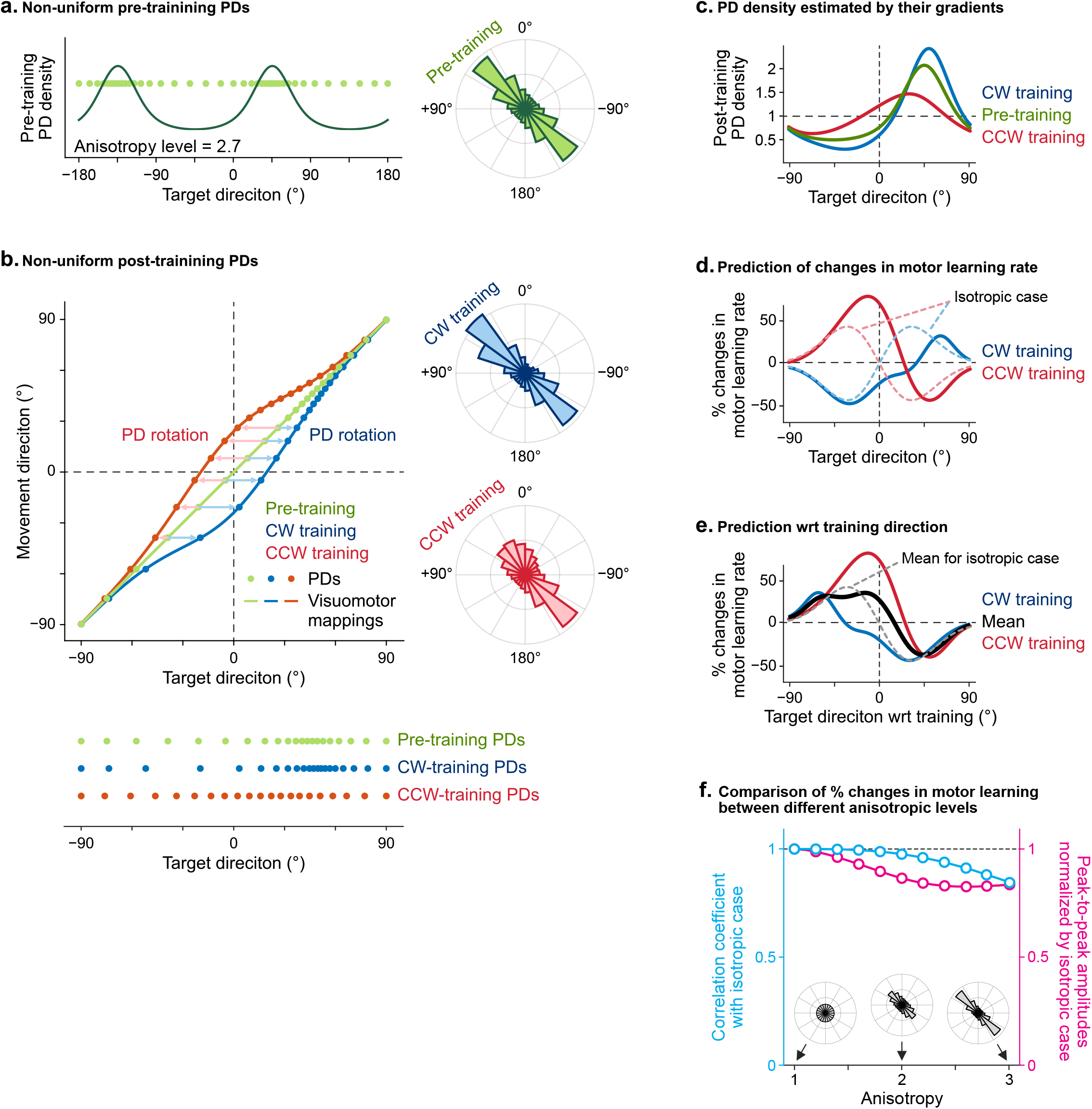
In the main text, we presented a simplified model where the baseline PD distribution was uniform (Figs. 1 and 2). However, the PD distribution in M1 has been shown to be non-uniform in monkeys, with a pattern that mirrors the marked anisotropy of limb inertia (Scott et al., 2001). Specifically, they found a x2.7 increase in PD density for high-inertia movement directions primarily driven by shoulder joint motion, which rotates the entire arm and corresponds to the 2nd and 4th quadrants of the horizontal plane, compared to low-inertia directions primarily driven by elbow motion, which rotates only the forearm and corresponds to the 1st and 3rd quadrants. We, therefore, simulated a baseline PD distribution that was elliptical rather than circular when viewed to polar coordinates, with the major axis projecting into quadrants 2 and 4, the minor axis projecting into quadrants 1 and 3, and an anisotropy ratio of 2.7. This distribution was fed into the model illustrated in Figures 1 and 2, and then, like in Figures 1 and 2, we examined the changes in (b-c) PD density and (d-e) learning rates predicted by the model for CW and CCW VMR training. For the uniform baseline PD distribution the changes induced by CW and CCW VMR training were identical when aligned to the direction of the training, but with an anisotropic baseline this is not the case, and so here we display the results separately for CW (blue) and CCW (red) training in panels b-e. Note that panel e shows target direction with respect to training direction as in Figures 2 and 4, which aligns model-predicted increases in density at PD rotation destinations and decreases in density at PD rotation origins. Note despite the marked anisotropy of the baseline distribution, the biphasic patterns of changes in learning rates are similar to the isotropic baseline case, with fairly subtle quantitative differences. Panel f presents an overall comparison between the patterns of training-aligned learning rate changes averaged across CW & CCW training for isotropic vs anisotropic baselines. Results are shown for anisotropy levels between 0 and 3, rather than only for the 2.7 level illustrated in b-e, for comparisons of both the shape (correlation coefficient, cyan) and the peak-to-peak amplitude (biphasic amplitude, pink) of the pattern of learning rate changes.

**Supplementary Data Figure 2.**
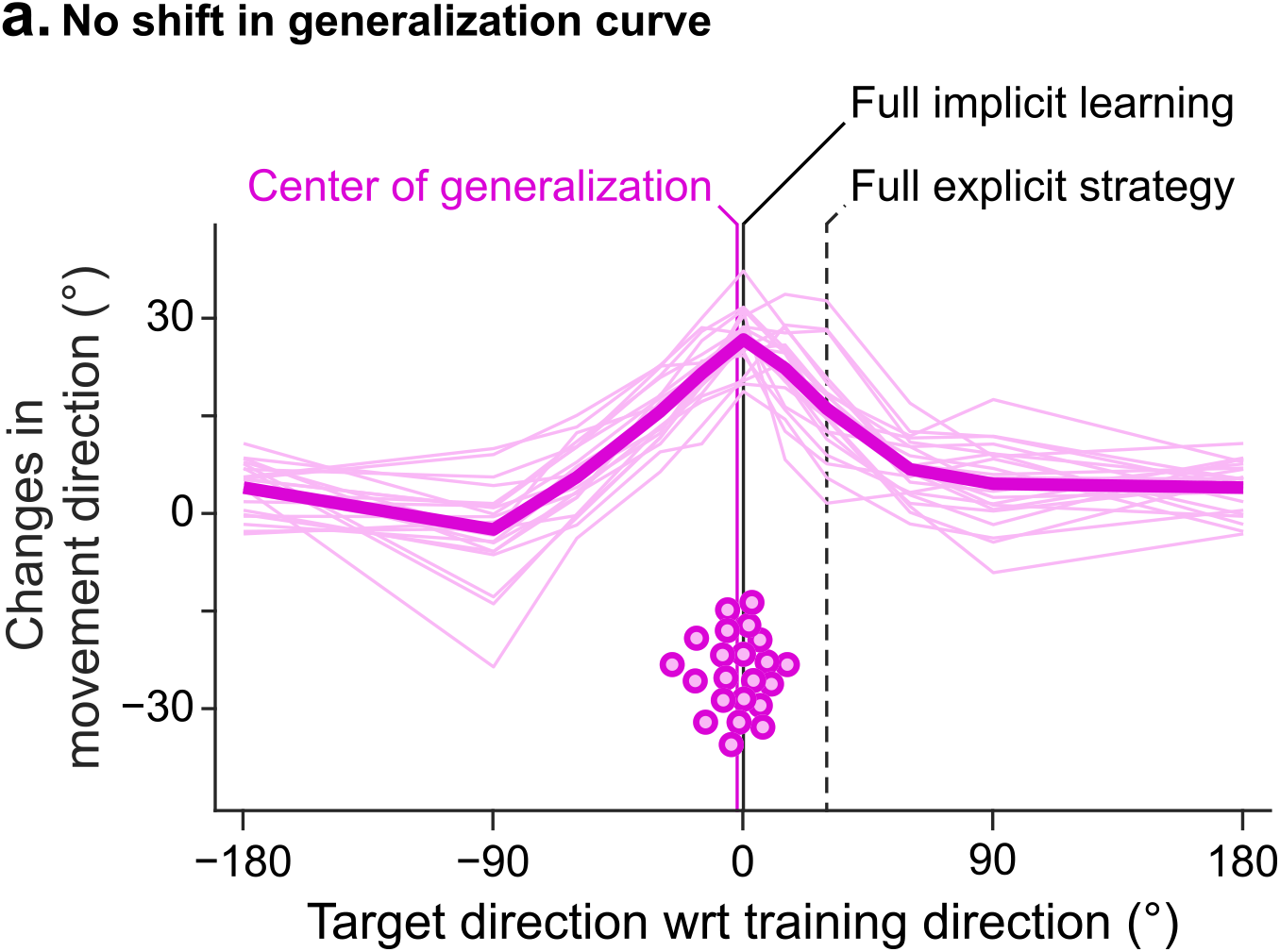
The VMR experiment we performed was designed to promote implicit learning with a gradual onset perturbation that was limited to 30°, low latency cursor feedback, the use of point-to-point rather than shooting movements, and the absence of aiming instructions (Brudner et al., 2016; Hadjiosif et al., 2024; Krakauer et al., 2019; Modchalingam et al., 2023; Neville and Cressman, 2018; Schween and Hegele, 2017). We therefore expected minimal use of explicit strategies. To check the extent to which this was the case, we made use of the finding that VMR learning is generalized around the aiming direction rather than the target direction (Day et al., 2016). Thus, the displacement between the center of the generalization curve and the target direction would indicate the amount of explicit aiming strategy. For the 30° VMR training we presented, the generalization curve would thus be centered at 30° if the sensorimotor learning relied on fully on explicit strategy, but 0° if it relied on fully on implicit learning. These generalization curves are plotted for all individual participants (thin lines) and for the mean data (thick line), with the generalization direction aligned to the training direction in both cases so that the explicit strategy predictions for CW and CCW training would match. To estimate the center for each individual’s generalization curve, we fit a gaussian function with a center shift (*θ*_*cs*_): *G*_*cs*_(Δθ) = λexp{−(*θ*_*cs*_ − Δ*θ*)^2^/2σ^2^}. We found that the center shift was −2.27° ± 2.15° (mean ± SEM), significantly different from the 30° full explicit strategy prediction (t(21) = 15.04, p = 1.02 × 10^−12^) but not different from the 0° full implicit learning prediction (t(21) = 1.06, p = 0.30). These results suggest little explicit contribution to the trained VMR learning.

